# Comparison of dimensionality reduction and feature selection for cognitive task decoding in functional magnetic resonance imaging

**DOI:** 10.1101/2025.10.27.684660

**Authors:** Corey J. Richier, Kyle A. Baacke, Sarah M. Olshan, Wendy Heller

**Author notes:** Correspondence (co-first authorship) concerning this article should be addressed to Corey J. Richier, 603 E Daniel St, Champaign, IL, 61820, or Kyle A. Baacke, 2441 E. Hartford Ave., Milwaukee, WI 53211. These authors contributed equally to the work (Corey J. Richier, Kyle A. Baacke).

## Abstract

**Background:** Advances in functional magnetic resonance imaging (fMRI) have led to the ability to study the brain across many contexts. However, the large number of features generated by functional connectivity approaches may overfit the data. These problems can be overcome with either feature selection (FS) or dimensionality reduction (DR), which can be applied to less complex models. We utilize two open-source datasets to compare the performance of DR/FS methods on cognitive task decoding using a suite of ML classifiers.

**New Method:** While DR and FS methods have been used previously in decoding research, no systematic comparison of their performance has been undertaken. Here, we compare available methods using commonly utilized machine learning libraries to establish which methods provide the best predictive performance. We then conduct statistical tests to examine the relative contributions of DR and FS methods and classifiers on decoding accuracy.

**Results:** Neither DR or FS was found to be superior. However, differences were identified across datasets and tasks. In the majority of methods and datasets, a peak in predictive performance was found using a small percentage of the total number of original features.

**Comparison with existing methods:** Some methods perform better than the baseline method of prediction with all available features or selecting features randomly. Decoding performance utilizing the HCP datasets with certain DR/FS methods exceeds that of deep learning approaches.

**Conclusions:** Simple machine learning models with DR/FS have competitive decoding performance. These results suggest a “sweet spot” for the tradeoff between the retention of features and predictive accuracy.

## 1. Introduction

Advances in the analysis of functional magnetic resonance imaging (fMRI) data have led to the ability to densely sample the human brain during a variety of contexts. Functional connectivity (FC) methods have shown promise for establishing linkages between brain function, behaviors, and cognition (Rogers et al., 2007). However, the large number of features generated by FC approaches (on the order of 10^3 to 10^7; Brown & Hamarneh, 2016) results in high dimensionality (known colloquially as the “curse of dimensionality”; Mwangi et al., 2014). A large number of features is undesirable due to the fact that the predictor variables can have high multicollinearity, thus reducing the interpretability of any given predictor (Alin, 2010). In predictive modeling contexts, a large number of features that exceed the number of observations may lead to models overfitting the data (Hua et al., 2009; Whelan & Garavan, 2014). This problem is generally overcome by employing either feature selection (FS) or dimensionality reduction (DR) methods. FS methods reduce the number of features by selecting particularly important or representative variables (Hauskrecht et al., 2007; Li et al., 2018), whereas DR methods transform existing variables into fewer variables that capture the original data’s variance (van der Maaten et al., 2009).

Arguably, one of the most foundational goals of neuroscience is examining how patterns of neural activity relate to mental processes. Decoding approaches have become particularly useful for examining relationships between cognitive states and representations of neural activity associated with those states (Poldrack, 2011), as they can allow for reverse inference (i.e., what is the behavioral state associated with a given pattern of neural activity). Decoding is flexible in its ability to be applied at various scales, from the level of cellular activity (Glaser et al., 2020) to whole brain recordings (Haynes & Rees, 2006), and is applicable for basic science (Haxby et al., 2001; LaConte, 2011) and clinical research (Rashid & Calhoun, 2020). Additionally, such methods have implications for brain computer interfaces (Rao, 2019). Recent initiatives in decoding have sought to make models as flexible as possible (Rubin et al., 2017) due to the fact that the nature of mental faculties are highly dimensional. Humans experience different stimuli and recruit a variety of different cognitive mechanisms to navigate the world. Effective decoding schemes therefore should be able to distinguish between as many different contexts and conditions as possible. Decoding schemes can be considered multi-class classification problems, in which the goal is to correctly classify the task associated with accompanying neural data. Given a large number of features in neural decoding approaches, methods to reduce the dimensionality of the input data may make these models more tractable. Thus, DR and FS approaches are suitable tools to improve classification accuracy.

Some DR methods and FS methods have been previously applied to decoding analyses (Mwangi et al., 2014). Among FS methods, random forest-based methods for feature selection have been found to outperform univariate methods (such as F-test score) for decoding approaches such as an object recognition task (Genuer et al., 2010). For DR methods, principal component analysis (PCA) has been found to be effective for decoding hand trajectory using electroencephalography and found it to be a more efficient method compared to using the mean value for all regions of interest at a given time point (Srisrisawang & Müller-Putz, 2022). DR and FS have been more formally compared in non-neuroimaging datasets. Anowar et al. (2021) compared multiple DR and FS methods across three different types of data modalities. In an electrocardiogram dataset, kernel PCA was found to perform best. However, in a mass spectroscopy dataset, it was found that laplacian eigenmaps were found to be the optimal method. Lastly, in a dataset for decoding mobility status from smartphone data, latent discriminant analysis was found to perform best. This emphasizes the need to explore the merits of different DR and FS methods in specific contexts, as the best-performing method depends on the structure of the input data.

Although deep learning methods can be considered the state of the art for machine learning approaches in neuroscience, they are not always the most advantageous method, depending on the circumstances. Linear models, by merit of their structure, can offer advantages with respect to interpretability (Varoquax et al., 2017). Deep learning models are often critiqued for being “black box”, because it is not straightforward to interpret which features are used for classification.

Knowing which brain features are implicated in certain cognitive processes, emotions, or disease states contributes not only to understanding basic mechanisms but has implications for applied utility (e.g., in the case of clinical disorders). Furthermore, more complex models like deep learning do not always perform better in supervised learning contexts, where simpler classifiers can perform comparably (Rudin, 2019). While there have been great large-scale advances in deep learning (for example, commercial LLMs), there have been criticisms of the environmental impact of large-scale models (Selvan et al., 2022). Some machine learning researchers have advocated that the efficiency of models is also a valid performance metric (Schwartz et al., 2020). While more classic machine learning models are less complex, they are more computationally efficient than some of the latest large-scale architectures. Thus, it can be easier to run many simpler models at scale to test hyperparameters or research questions when many permutations of the data or models must be examined. Additionally, the scaling of deep learning research has resulted in certain institutions, including private corporations, having access to methods that are simply intractable for many researchers (Besiroglu et al., 2024; Shu et al., 2024). A lack of access to computational resources that can employ such large-scale deep learning models on imaging data can lead to scientific and research inequity (Nielsen & Andersen, 2021). Thus, determining inclusive machine learning methods for neuroscience, which operates using highly dimensional data, will be important for the continued participation of researchers around the globe, who may not always have access to state of the art computational resources (Ahmed & Wahed, 2020; Probst, 2023).

While DR and FS have been used in analyses of fMRI data, no formal comparisons have taken place to examine their efficacy in a decoding task. Taking inspiration from previous research in other fields (Anowar et al., 2021), the present analysis sought to compare numerous DR and FS methods on task-based fMRI for how well they may improve classification accuracy. We compared decoding accuracy of cognitive tasks in multiple datasets using various DR and FS methods. We used seven DR/FS methods to reduce the input space generated by functional connectivity values from parcellated fMRI sessions in two publicly available datasets. We then used classifier models to decode tasks from the resulting input feature sets. The cross-validation accuracy of the resulting models was then used to evaluate their efficacy. Stepwise hierarchical regressions were then used to evaluate whether the classifier, DR/FS method, and interactions between classifier and method impacted model performance after accounting for the impact of the number of features used in each model.

## 2. Method

### 2.1 Datasets

Data used in the present work were obtained from the Human Connectome Project (HCP) database (Van Essen et al., 2012; https://db.humanconnectome.org/) and the UCLA Consortium for Neuropsychiatric Phenomics dataset (Poldrack et al., 2016; obtained from https://openneuro.org/datasets/ds000030, version 1.0). Data from the HCP project were preprocessed using the HCPPipelines BIDS App wrapper (https://github.com/BIDS-Apps/HCPPipelines; Glasser et al., 2013; Smith, 2018). Preprocessing for the UCLA dataset relied on fMRIPrep 20.2.0 (Esteban et al., 2019, 2023), which is based on Nipype 1.5.1 (Esteban et al., 2022; Gorgolewski et al., 2011); RRID:SCR_002502). Further details about preprocessing methods are provided in Supplementary Materials.

In HCP, the subjects completed tasks as follows: 1) a working memory task, as measured via an N-back paradigm, 2) a motor task, where participants moved their fingers, toes, or tongue, 3) a language processing task, where participants completed basic arithmetic and story comprehension questions, 4) a social cognition tasks, where participants were to infer theory of mind in a series of inanimate objects moving in a way that suggested interaction, 5) a relational processing task, where they were presented with different images and told to compare and contrast the characteristics of these images, 6) an emotional processing task, where they were asked to compare facial expressions across images and indicated whether a series of faces were the same or not, and 7) an incentive processing task, where they were presented with a risk/reward paradigm with potential to gain or lose money through a series of decisions. In the UCLA study, participants engaged in a 1) balloon analog risk task, where participants had to risk maximizing the amount of money acquired while risking losing what they had earned, a paired association memory task, with 2) an encoding phase and 3) a retrieval phase, 4) a spatial working memory task, 5) a stop signal task, where participants underwent a go no-go paradigm, 6) a task-switching task, where participants were asked to respond to different stimuli in changing sequences, 7) a breath-holding task, and 8) a resting state scan with eyes open. Further details of these tasks can be obtained in Barch et al. (2013) for HCP and Poldrack et al. (2016) for UCLA.

### 2.2 Data processing

The non-smoothed task-fMRI data for each participant were parcellated using the NiftiLabelsMasker function in NiLearn v0.9.0 (Abraham et al., 2014; RRID:SCR_001362), referencing the 200 parcel 7 network atlas generated by Schaefer et al. (2018). The parcellation combines voxels that belong to specific brain regions. The parcellated timeseries were then concatenated for each subject for each task (RL and LR phase encoding). The mean of all voxel BOLD signals in each parcel was then subtracted from the values in the timeseries to center values for each parcel around 0. These values in each parcel were then Fisher-z transformed. After this, every parcel was correlated with one another using the Pearson correlation to generate functional connectivity matrices for each task session for each subject. The unique values from this matrix (i.e., the lower or upper diagonal) were extracted to serve as input data for the classifiers. This process was repeated for every subject’s available task data. Thus, the total number of observations from each data set represented the total number of unique task runs across subjects (such that each subject was represented in the data the number of times they completed a unique task). The final matrix was made up of columns representing the different connectivity values for each pair of parcels, and rows representing a unique task session. Each column of this data matrix was then standardized with a *z*- score transformation.

### 2.3 Feature selection and dimensionality reduction methods

All dimensionality reduction and prediction generation steps were conducted with Python 3.6.15. The present DR and FS methods were selected because they can be fitted to a training set of data and then fitted on a separate test set. The full feature set (19,900) was used to compare the various DR and FS as a baseline. For another baseline comparison, random feature sets were selected from the full feature set. For each of the feature set sizes generated using the other DR/FS methods, 10 sets of features were randomly selected from the initial 19,900 features.

#### 2.3.1 Feature selection methods

##### 2.3.1.1 Hierarchical Clustering

Hierarchical clustering (HC) was performed as follows using scipy v1.3.3 (Virtanen et al., 2020; RRID:SCR_008058). Spearman correlations between variables were used as an input euclidean distance matrix to perform Ward’s linkage to identify the hierarchical structure of clusters by minimizing variance within each cluster. The resulting linkage matrix was then used to form flat clusters for each within-cluster cophenetic distance cap between 1 and 250. Unique cluster structures were saved for use in the prediction generation phase.

##### 2.3.1.2 Permutation Importance

Permutation importance methods constitute a broad class of techniques that randomly shuffle feature values to disrupt any systematic association between predictors and the outcome, using this permuted data to generate a null distribution of test statistics or *p*-values for comparison(Winkler et al., 2014). The permutation importance method employed in scikit-learn 0.23.2 ((Pedregosa et al., 2011); RRID:SCR_002577) permutes each variable in the dataset and evaluates the change in overall accuracy when the feature is permuted. Features that have a greater reduction in accuracy when they are permuted are deemed to be more important for successful prediction.

##### 2.3.1.3 Select from model

The select from model (SFM) method retrieves features from the model training process based on different importance metrics. For the random forest, the Gini impurity decrease is the metric used for establishing feature importance. In the support vector machine, model coefficients are selected by their magnitude. SFM was performed using scikit-learn v0.23.2.

#### 2.3.2 Dimensionality reduction methods

##### 2.3.2.1 Principal Component Analysis

Principal component analysis (PCA) is a commonly used dimensionality reduction method that identifies orthogonal axes in the data, known as principal components. These axes capture the maximal amount of variance in the data. PCA projects data onto these axes, and these are used as new predictor variables. Therefore, fewer variables can be used to capture a greater amount of variation in the data. All components were retained for prediction generation. For all feature set sizes generated using hierarchical clustering, the same number of components was retained as input in the prediction generation step. PCA was performed using scikit-learn v0.23.2.

##### 2.3.2.2 Kernel Principal Component Analysis

Kernel PCA (kPCA) is a similar method to PCA that projects data to higher dimensional space to better account for potential non-linear relationships in the data. The present analysis utilized kPCA with linear and radial basis function kernels. kPCA was performed using scikit-learn 0.23.2.

##### 2.3.2.3 Truncated Singular Value Decomposition

Singular value decomposition (SVD) is a dimensionality reduction method that decomposes any matrix into three matrices that contain information about variation in the data. In truncated SVD (tSVD) only a specified number of singular values and singular vectors, which can be used as new predictor variables, are retained. tSVD was performed using scikit-learn 0.23.2.

##### 2.3.2.4 Linear Discriminant Analysis

Linear discriminant analysis (LDA) is a supervised learning method utilizing DR to find the optimal linear combination between components of the predictor. LDA will generate an equal number of predictor variables to the number of classes to predict. These variables attempt to maximize the variation between classes while minimizing the variation within the classes.

#### 2.3.3 Classifiers

Research has demonstrated that the choice of model can have an effect on performance in machine learning decoding analyses with fMRI data (Jollans et al., 2019). Thus, the present study utilized three types of models with different architectures. A random forest classifier (RFC), a support vector machine (SVM), and ridge regression (RR) were utilized. All three models were given data from all of the aforementioned DR and FS methods. A grid search was utilized to examine the best combination of hyperparameters for each of the different classifiers. For the SVM, all models were instantiated with a linear kernel, a squared hinge loss function, and 100,000 max iterations.

Values of C were iterated over in the grid search (0.1, 1, 10). For the RFC, the number of estimators were grid searched over 100 and 200, and the minimal samples for leafing were set at 2 and 5. For the RR, alphas were set to .001, .01, .1, 1, and 10.

##### 2.3.4 Training and testing procedure

Ninety percent of the data was initially selected for the training procedure. A group k-fold method was used such that no subjects appeared in both the training and testing data, as this might result in inflated estimates of testing set accuracy (Varoquax et al., 2017). This procedure entailed a further 90/10 split ten-fold cross validation to obtain the best features for predictive accuracy. These features were then applied to the held out 10% of the data and evaluated for accuracy. Each classifier was cross validated with each feature set size and DR/FS. A group k-fold with five splits was used for training (roughly 18% was used as the testing data).

## 3. Results

### 3.1 Performance by dataset, classifier and DR/FS method

Figure 1 demonstrates a performance curve of each method in the HCP dataset, and Figure 2 demonstrates the same in the UCLA dataset. The curves were generated by comparing each method to selecting features at random of the same distribution of features sizes (shown as grey dots) to serve as a benchmark for each method. Several trends were evident based on the figure. Many methods were at their highest classification accuracy between the order of 10-1,000 features. These methods seem to taper with respect to classification performance when the number of features becomes large enough. However, some methods did not perform well at any size. Performance of the SVM in the HCP dataset was poor over most of the possible feature sizes, except for a larger number of features in SFM, RFPI, and HC. For tSVD, PCA, and kPCA, most numbers of features result in a very low degree of accuracy, suggesting that these methods did not capture meaningful task-related variance in the data. However, this effect was not noted in the UCLA dataset, in which these methods are substantively more effective in classification of tasks than randomly selecting features. Additionally, in the HCP dataset, hierarchical clustering did not perform better than randomly selecting features. This effect was also noted in the UCLA dataset.

**Figure 1.**
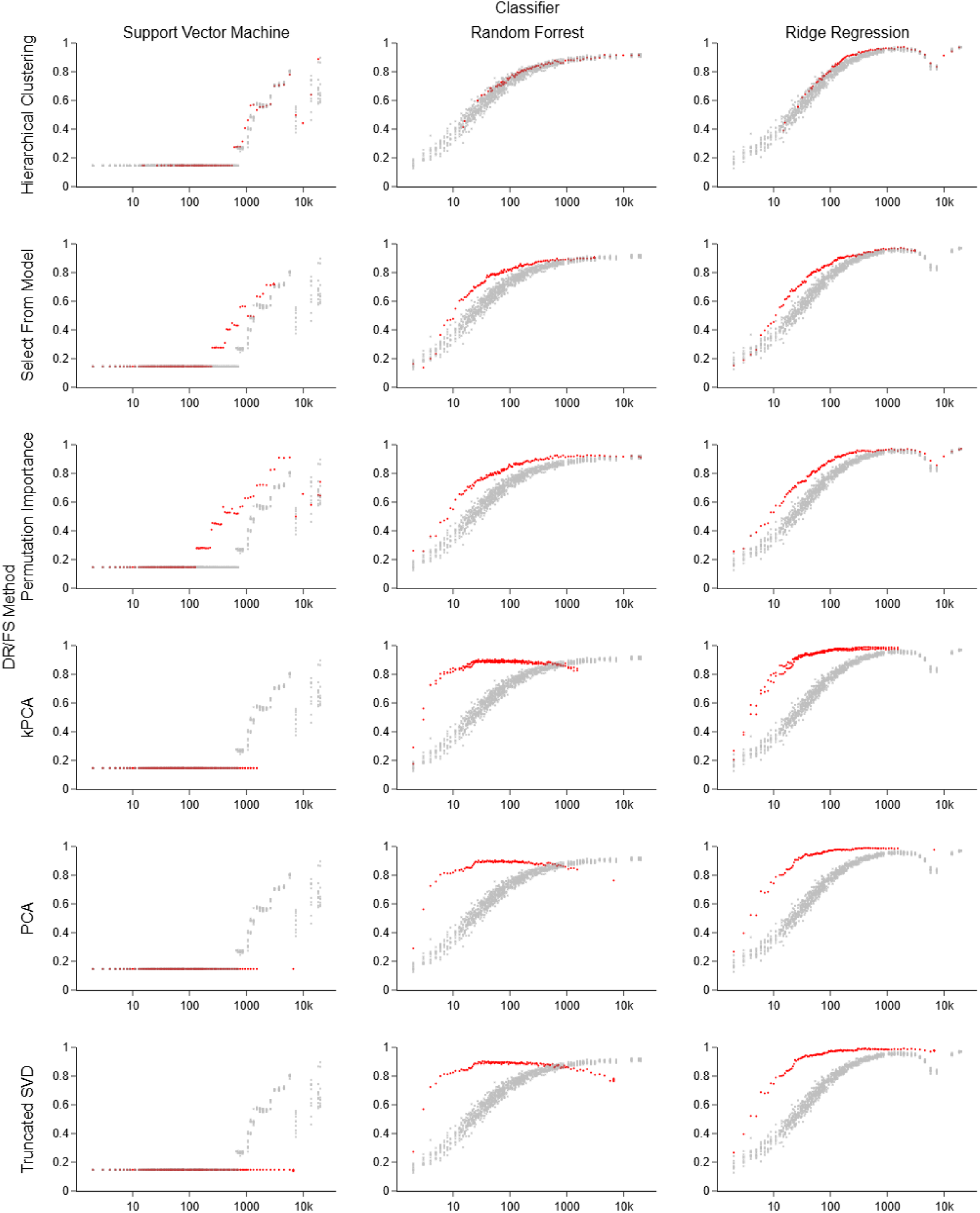
Performance Curve of Each DR and FS Method Across Feature Sizes in HCP. *Note:* Red dots represent accuracy of that classifier at a certain number of features. Grey dots are randomly selected features to serve as a comparison to the method. LDA is not displayed because only one set of features was utilized. Numbers in parentheses represent how many observations were used to train the model. RFPI: Random Forest Permutation Importance, tSVD: Truncated Singular Value Decomposition, PCA: Principal Component Analysis, kPCA: Kernel Principal Component Analysis

**Figure 2.**
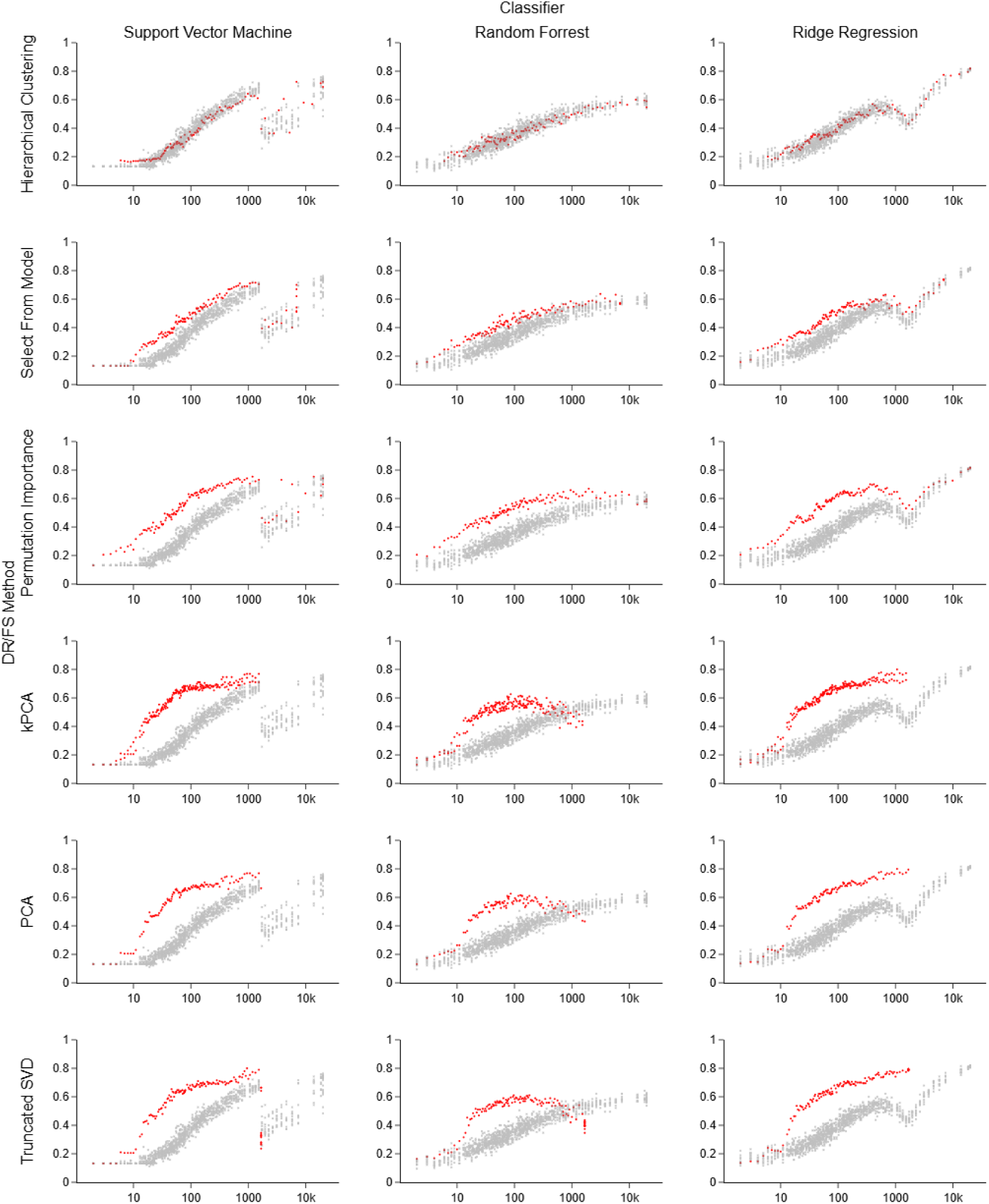
Performance Curve of Each DR and FS Method Across Feature Sizes in UCLA. *Note:* Red dots represent accuracy of that classifier at a certain number of features. Grey dots are randomly selected features to serve as a comparison to the method. LDA is not displayed because only one set of features was utilized. Numbers in parentheses represent how many observations were used to train the model. RFPI: Random Forest Permutation Importance, tSVD: Truncated Singular Value Decomposition, PCA: Principal Component Analysis, kPCA: Kernel Principal Component Analysis

### 3.2 Best performing classifier and DR/FS method by dataset

Table 1 reports the best performing classifier for each dataset. Across all methods and feature sizes, utilizing ridge regression with 1,959 features derived from tSVD was the most accurate classification method in the HCP dataset. In UCLA, the most effective method for ridge regression used all 19,900 features, whereas the second most effective method that reduced the number of features was again tSVD using a support vector machine.

**Table 1.** Best accuracy of feature selection/dimensionality reduction method by classifier and dataset.

| Classifier | HCP<br>(Number of features - DR/FS Method) | UCLA (Number of features<br>- DR/FS Method) |
| --- | --- | --- |
| Support Vector Machine | 91.18% (5,774 - RFPI) | 80.00% (948 - tSVD) |
| Random Forest Classifier | 96.66% (7 - LDA) | 67.89% (8 - LDA) |
| Ridge Regression | 99.06% (1,959 - tSVD) | 82.11% (19,900 - Hierarchical) |

*Note:* HC Hierarchical Clustering, RFPI: Random Forest Permutation Importance, SFM: Select From Model, tSVD: Truncated Singular Value Decomposition, PCA: Principal Component Analysis, kPCA: Kernel Principal Component Analysis, LDA: Linear Discriminant Analysis

Table 2 lists the overall accuracy of each type of DR/FS method. Of note, ridge regression was found to be the method which gave rise to the most accurate DR/FS results for each method. In HCP, the most accurate method was kPCA, although most methods were closely accurate (roughly 97-98%). In UCLA, the two most accurate methods were selected using the entire feature set or close to it (HC and RFPI). The next most accurate methods that did not default to the entire (or close to) the entire feature set were kPCA, PCA, and tSVD, which were 80% accurate and used 948 features for tSVD, and 1,065 features for PCA and kPCA. Of note, these results do not suggest an obvious performative edge in DR or FS methods in either dataset.

**Table 2.** Overall accuracy of feature selection/dimensionality reduction method by dataset across all feature.

| DR/FS Method | HCP (Number of features - Classifier) | UCLA (Number of features - Classifier) |
| --- | --- | --- |
| HC | 97.19% (1,958 - RR) | 82.11% (19,900 - RR) |
| kPCA | 98.935 (480 - RR) | 80.00% (1,065 - RR) |
| LDA | 97.06% (7 - RR) | 73.68% (8 - RR) |
| PCA | 98.93% (419 - RR) | 80.00% (1,065 - RR) |
| RFPI | 97.19% (1,185 - RR) | 81.58% (19,878 - RR) |
| SFM | 97.33 (1,728 - RR) | 73.68% (6,885 - RR) |
| tSVD | 90.06% (1,959 - RR) | 80.00% (948 - RR) |

*Note:* HC Hierarchical Clustering, RFPI: Random Forest Permutation Importance, SFM: Select From Model, tSVD: Truncated Singular Value Decomposition, PCA: Principal Component Analysis, kPCA: Kernel Principal Component Analysis, LDA: Linear Discriminant Analysis, RR: Ridge Regression

To examine overall trends in the classification accuracy, we chose to examine the top 10% highest testing accuracy DR/FS methods for each dataset. Overall, the average testing accuracy in the top 10% of the HCP dataset was 96.24% (SD = 1.53%, range = 93.58-99.06%) and 71.79% (SD = 4.07%, range = 66.84-82.11%) in the UCLA dataset, indicating that decoding was more accurate in the HCP dataset. The results of the top 10% best performing classifiers with respect to decoding accuracy are reported in table 3. The best performing overall classifier was ridge regression. Random forest and ridge regression performed better in HCP than UCLA; however, the support vector machine performed better in the UCLA dataset.

**Table 3.**
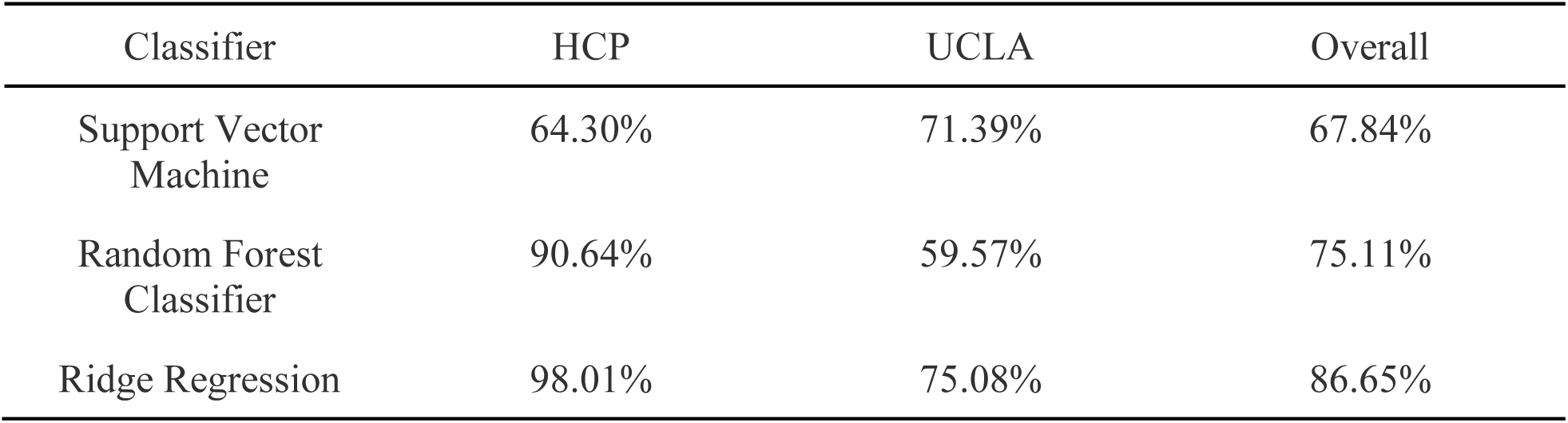
Decoding accuracy by classifier in top 10% best performing DR/FS methods for each classifier.

Next, the top 10% of best performing DR/FS methods were examined. These results are shown in table 4. In addition to the individual DR/FS methods, both the average of all three classifiers’ accuracy using the full set of features (i.e., no DR/FS) and randomly selected features are reported as a baseline comparison. In HCP, all methods performed better than the random selection or full feature set. In the UCLA dataset, the full feature set performed better than SFM and HC, and HC was comparable to random feature selection. Across both datasets, tSVD was the best performing DR method, whereas RFPI was the best performing FS method.

**Table 4.**
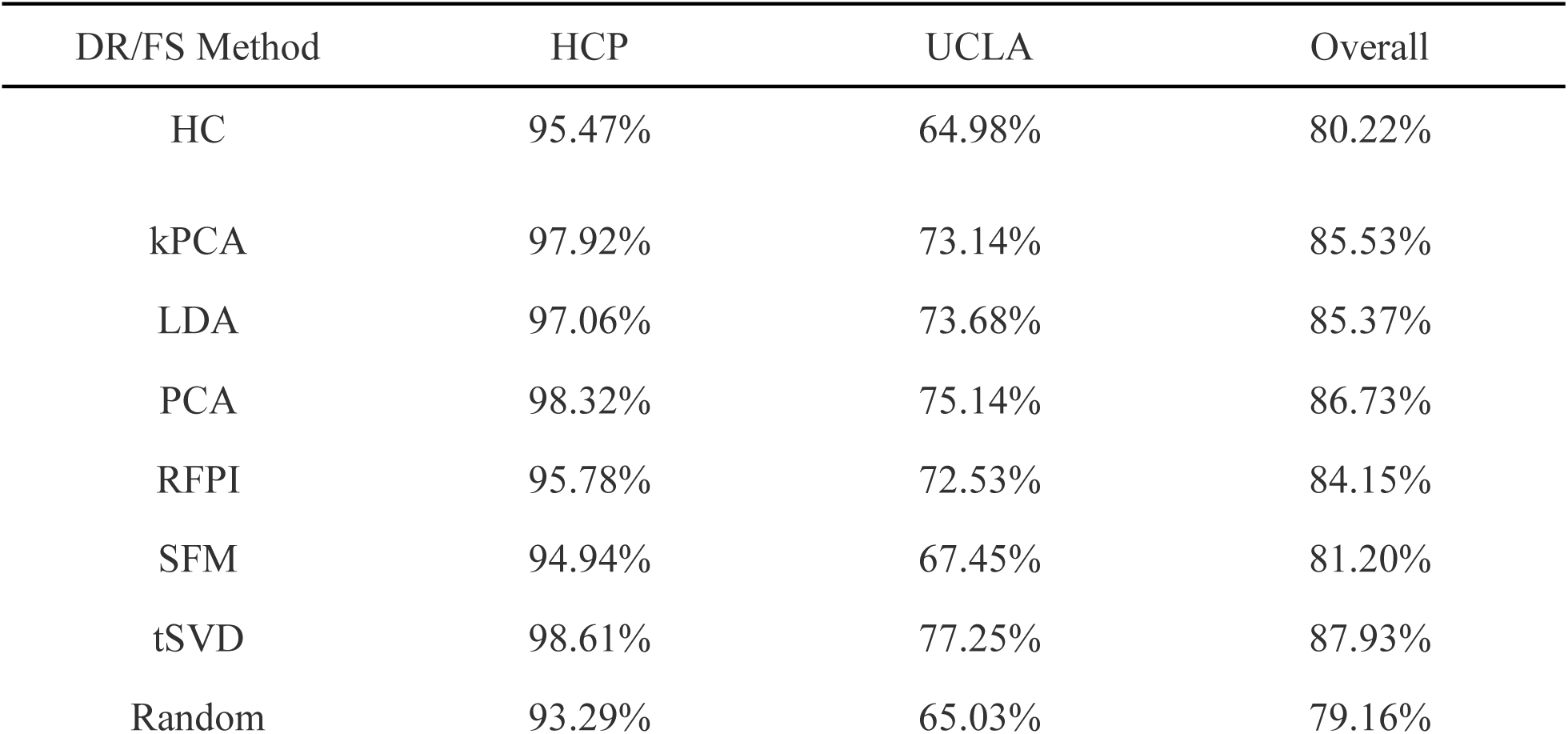
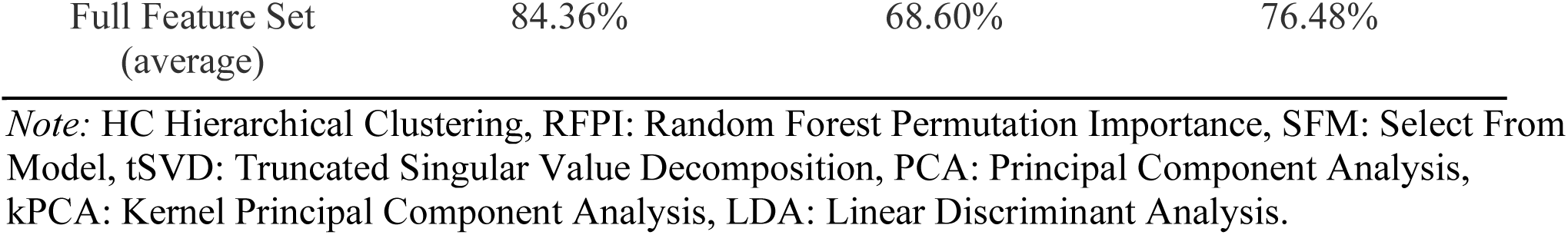
Overall accuracy of feature selection/dimensionality reduction method by dataset across all feature sizes.

### 3.3 Task-level results

In addition to overall performance across DR/FS methods and classifiers, we examined differences in decoding accuracy across specific tasks in the top 10% of DR/FS sets (Tables 5 and 6). It is worth noting that for the HCP dataset, the support vector machine struggled to classify several tasks, such as the emotion, gambling, and working memory tasks. This explains the relatively lower degree of decoding accuracy seen in the SVM relative to other models. The social processing and relational task was the most accurately classified. In the UCLA dataset, the breath holding task was the most accurately classified, whereas the task switching task was the least accurately classified.

**Table 5.** Accuracy of prediction of each class in the HCP Dataset by classifier in top 10% decoding accuracy.

| Task | Support Vector<br>Machine Mean<br>Accuracy | Random Forest<br>Mean Accuracy | Ridge Regression<br>Mean Accuracy | Overall Mean<br>Accuracy |
| --- | --- | --- | --- | --- |
| Emotion Task | 34.14% | 88.06% | 98.49% | 73.56% |
| Gambling Task | 18.96% | 82.82% | 95.97% | 65.92% |
| Language Task | 89.25% | 95.54% | 99.28% | 94.69% |
| Motor Task | 87.01% | 98.27% | 99.21% | 94.83% |
| Relational Task | 87.08% | 82.47% | 95.25% | 88.27% |
| Social Processing<br>Task | 93.54% | 96.48% | 99.61% | 96.54% |
| Working Memory<br>Task | 43.08% | 90.80% | 98.27% | 77.39% |

**Table 6.** Accuracy of prediction of each class in the UCLA Dataset by classifier.

| Task | Support Vector<br>Machine Mean<br>Accuracy | Random Forest<br>Mean Accuracy | Ridge Regression<br>Mean Accuracy | Overall Mean<br>Accuracy |
| --- | --- | --- | --- | --- |
| Spatial Working<br>Memory | 69.25% | 54.88% | 73.04% | 65.72% |
| Balloon Analog Risk | 64.54% | 53.88% | 67.98% | 62.13% |
| Paired Association<br>Memory (Recall) | 71.73% | 67.51% | 75.87% | 71.70% |
| Paired Association<br>Memory (Encoding) | 65.40% | 52.91% | 63.98% | 60.76% |
| Task Switch | 59.32% | 50.96% | 66.45% | 58.91% |
| Stop Signal | 73.36% | 56.86% | 78.33% | 69.52% |
| Rest | 82.08% | 65.99% | 86.29% | 78.12% |
| Breath Holding | 83.91% | 74.31% | 85.65% | 81.29% |

Tables 7 and 8 examine the relative contributions of each DR and FS method to overall task decoding accuracy in the top 10% of decoding accuracy for each method. The overall task level decoding accuracy in HCP was quite high, and comparable to using the full feature sets. In UCLA, some DR/FS methods were not able to achieve greater classification accuracy in some of the tasks relative to using the full feature set, or the top performing groups of randomly selected features.

**Table 7.** Accuracy of prediction of each class in the HCP Dataset by DR/FS method.

| Task | HC | LDA | PCA | RFPI | tSVD | kPCA | SFM | Random | Full<br>Feature<br>Set |
| --- | --- | --- | --- | --- | --- | --- | --- | --- | --- |
| Emotion | 96.04% | 98.11% | 98.74% | 95.49% | 98.80% | 98.48% | 92.96% | 92.92% | 97.17% |
| Gambling | 90.59% | 92.73% | 96.76% | 90.88% | 97.52% | 95.63% | 90.93% | 87.91% | 93.64% |
| Language | 97.19% | 99.04% | 99.61% | 96.31% | 99.76% | 99.01% | 97.69% | 96.18% | 99.04% |
| Motor | 96.61% | 97.25% | 99.58% | 98.74% | 99.59% | 99.24% | 96.97% | 96.59% | 97.25% |
| Relational | 94.21% | 94.17% | 95.57% | 94.02% | 96.33% | 95.08% | 93.47% | 90.11% | 95.15% |
| Social<br>Processing | 98.88% | 99.06% | 99.69% | 98.88% | 99.46% | 99.59% | 97.42% | 97.45% | 99.06% |
| Working<br>Memory | 94.93% | 99.09% | 98.28% | 96.22% | 98.77% | 98.40% | 95.23% | 91.98% | 99.09% |

*Note:* HC Hierarchical Clustering, RFPI: Random Forest Permutation Importance, SFM: Select From Model, tSVD: Truncated Singular Value Decomposition, PCA: Principal Component Analysis, kPCA: Kernel Principal Component Analysis, LDA: Linear Discriminant Analysis

**Table 8.** Accuracy of prediction of each class in the UCLA Dataset by DR/FS method.

| Task | HC | LDA | PCA | RFPI | tSVD | kPCA | SFM | Random | Full<br>Feature<br>Set |
| --- | --- | --- | --- | --- | --- | --- | --- | --- | --- |
| Spatial<br>Working<br>Memory | 66.20% | 73.08% | 73.30% | 70.06% | 73.73% | 71.21% | 66.19% | 63.41% | 80.77% |
| Balloon<br>Analog<br>Risk | 58.55% | 76.00% | 69.49% | 70.27% | 75.02% | 67.06% | 61.79% | 57.98% | 76.00% |
| Paired<br>Association<br>Memory<br>(Recall) | 64.48% | 72.22% | 75.87% | 74.62% | 78.59% | 72.38% | 70.03% | 65.91% | 83.33% |
| Paired<br>Association<br>Memory<br>(Encoding) | 57.41% | 61.11% | 61.11% | 71.02% | 62.74% | 60.26% | 61.40% | 61.13% | 72.22% |
| Task<br>Switch | 56.06% | 53.85% | 65.49% | 63.31% | 71.20% | 63.25% | 60.53% | 55.15% | 76.92% |
| Stop Signal | 66.43% | 73.08% | 74.40% | 74.74% | 78.52% | 76.55% | 69.64% | 66.64% | 84.62% |
| Rest | 70.16% | 84.62% | 89.12% | 78.27% | 89.87% | 83.39% | 75.40% | 71.56% | 92.31% |
| Breath<br>Holding | 78.30% | 92.00% | 88.57% | 78.27% | 84.49% | 87.11% | 73.58% | 77.75% | 84.00% |

*Note:* HC Hierarchical Clustering, RFPI: Random Forest Permutation Importance, SFM: Select From Model, tSVD: Truncated Singular Value Decomposition, PCA: Principal Component Analysis, kPCA: Kernel Principal Component Analysis, LDA: Linear Discriminant Analysis

Some tasks, such as the social processing task, had high classification accuracy regardless of method employed. The UCLA dataset overall had lower task-based decoding accuracy than HCP. The breath holding task in particular had quite discrepant accuracies with respect to different methods (i.e., 92.00% with LDA, but 73% with SFM).

### 3.5 Stepwise hierarchical regression

For each of the two datasets, a stepwise hierarchical regression was performed to evaluate the impact of feature set size, classifier type, DR/FS method, and interactions between DR/FS methods and classifier types on classifier performance. In the first step, only the number of features and three transformations of the number of features (log, square root, and cubed root) were included in the models. For the UCLA dataset, 54.29% of variance in model accuracy was accounted for by these four variables (*F*[4, 5911]=1757.44, *p* < 0.001). For the HCP dataset, 15.93% of variance in model accuracy was accounted for by these four variables (*F*[4, 5818]=276.77, *p* < 0.001). The second step added dummy coded variables to indicate which classifiers were used (with SVM being the implicit baseline). Adding factors as classifier resulted in a significant increase in variance accounted for in both the UCLA dataset (*F*[6, 5909]=1260.82, *p* < 0.001 *R^2^*= 0.56; Δ*R*^2^ = 0.02, *F*[2,5909] = 122.77, *p* <0.001) and HCP dataset (*F*[6, 5818]=276.77, *p* < 0.001 *R^2^* = 0.853; Δ*R*^2^ = 0.69, *F*[2,5816] = 13692, *p* <0.001). The third step added additional dummy coded variables to represent which DR/FS methods were used with the implicit baseline of randomly selected features. Adding predictors indicating DR/FS method further increased the amount of variance accounted for in model accuracy for both datasets (UCLA: *F*[12, 5903]= 2550.96, *p* < 0.001 *R^2^* = 0.84; Δ*R*^2^ = 0.27, *F*[6,5903] = 1685, *p* <0.001; HCP: *F*[12, 5810]= 3593.78, *p* < 0.001 *R^2^* = 0.88; Δ*R*^2^ = 0.03, *F*[6,5810] = 232, *p* <0.001). In the fourth and final step, interaction terms between classifier and method were included. Adding the interaction terms resulted in small but significant increases in *R*^2^ in both datasets (UCLA: *F*[24, 5891]= 1358.07, *p* < 0.001 *R^2^*= 0.85; Δ*R*^2^ = 0.008, *F*[12,5891] = 26.89, *p* <0.001; HCP: *F*[24, 5798]= 2333.05, *p* < 0.001 *R^2^*= 0.91; Δ*R*^2^ = 0.02, *F*[12,5798] = 128.2, *p* <0.001). Overall model fit indices are reported in table 9, and coefficients from the final models are reported in table 10.

**Table 9.** Stepwise Hierarchical Regression Overview.

| UCLA |  |  |  |  | HCP |  |  |  |
| --- | --- | --- | --- | --- | --- | --- | --- | --- |
| Model | $df$ | $F$ | $p$ | $R^2_{Adj}$ | $df$ | $F$ | $p$ | $R^2_{Adj}$ |
| $N$ Features | (4,5911) | 1757.44 | <.0001 | 0.543 | (4,5818) | 276.77 | <.0001 | 0.159 |
| $N$ Features + Classifier | (6,5909) | 1260.82 | <.0001 | 0.561 | (6,5816) | 5616.96 | <.0001 | 0.853 |
| $N$ Features + Classifier + Method | (12,5903) | 2550.96 | <.0001 | 0.838 | (12,5810) | 3593.78 | <.0001 | 0.881 |
| $N$ Features + Classifier + Method + Interaction Terms | (24,5891) | 1356.07 | <.0001 | 0.846 | (24,5798) | 2333.05 | <.0001 | 0.906 |

**Table 10.**
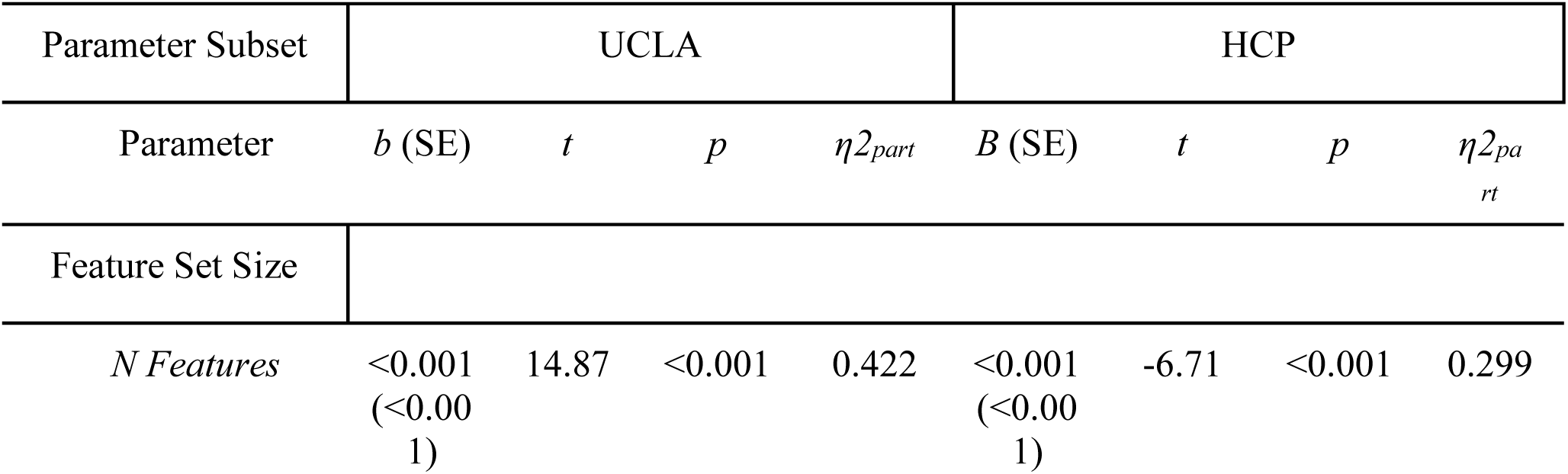
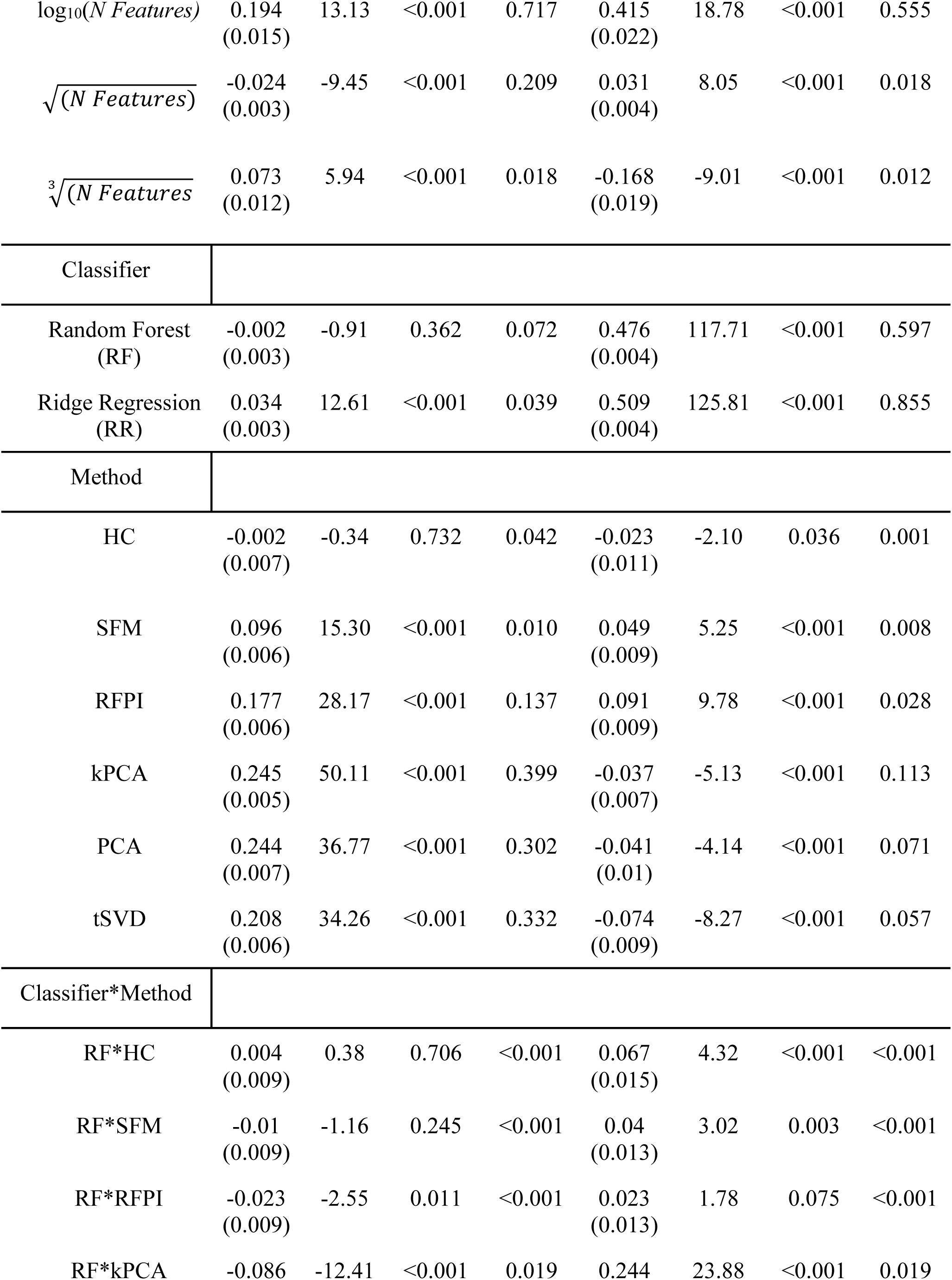
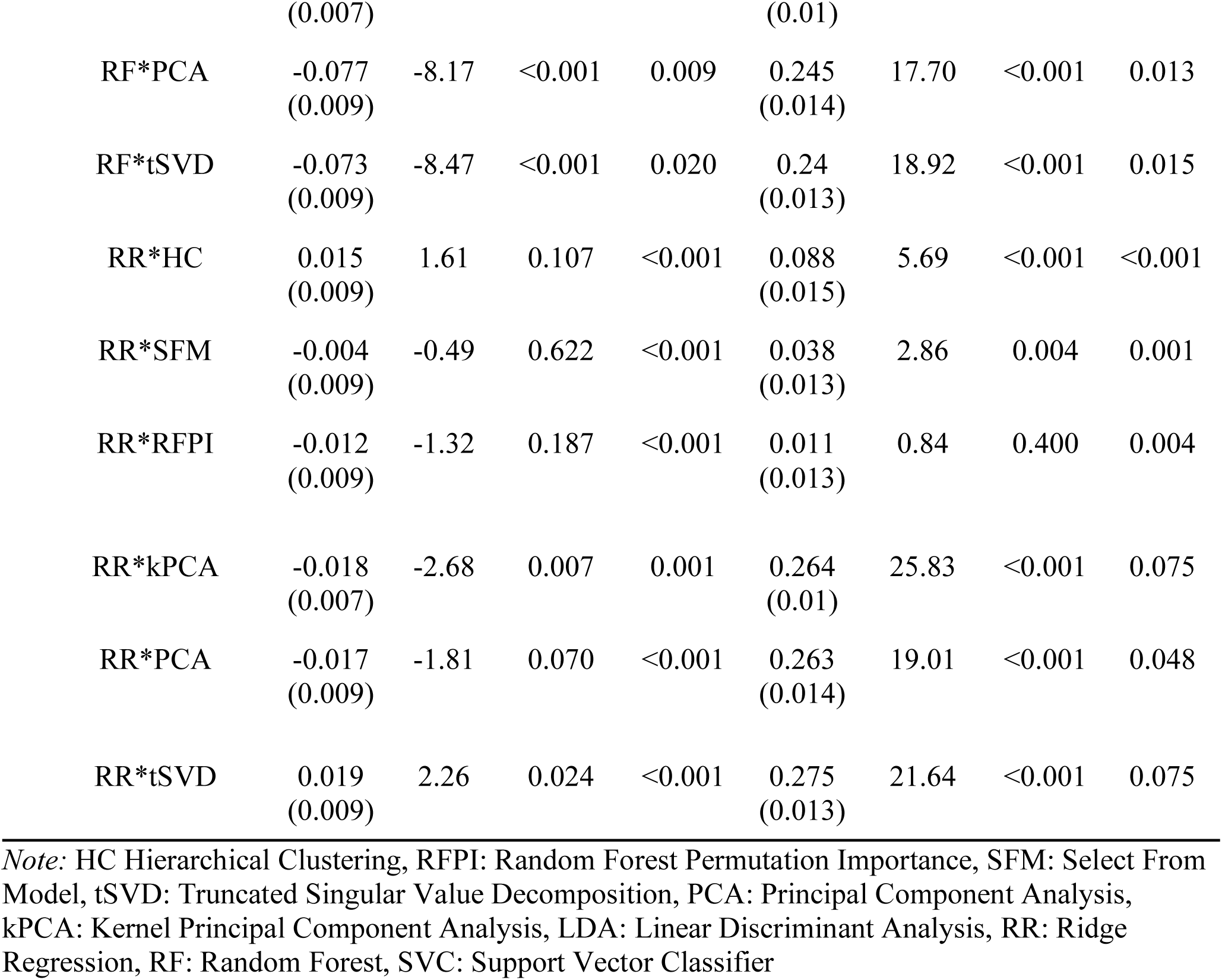
Final Model Regression Results.

Due to the exceptionally low performance of the SVMs in the HCP dataset, the entire stepwise regression procedure was repeated without SCM results for this dataset (Table 11). The results from the regressions in HCP without data from SVMs more closely mirrored the results from the UCLA dataset. In the first step, 66.2% of the variance in model accuracy was accounted for by the number of features and the three transformations thereof (*F*[4, 3877] = 1897.80, *p* < 0.001). The second step, which included only one dummy-coded variable to indicate whether RR was used instead of RF, accounted for a significantly greater portion of variance in overall model accuracy (*F*[5, 3876] = 1592.54, *p* < 0.001, *R^2^* = 0.672; Δ*R*^2^ = 0.01, *F*[1,3876] = 126.25, *p* <0.001). The addition of variables indicating which DR/FS method was used further improved model performance to a significant degree (*F*[11, 3870] = 2486.92, *p* < 0.001, *R^2^_Adj_* = 0.876; Δ*R*^2^ = 0.20, *F*[6,3870] = 1058.90, *p* <0.001). The final step, which included interaction terms between classifier and DR/FS method accounted for a small but statistically significant portion of variance in model accuracy (*F*[17, 3864] = 1621.99, *p* < 0.001, *R^2^* = 0.877; Δ*R*^2^ = 0.001, *F*[6, 3864] = 5.37, *p* <0.001).

**Table 11.** Stepwise Hierarchical Regression Overview.

| UCLA |  |  |  |  | HCP |  |  |  |
| --- | --- | --- | --- | --- | --- | --- | --- | --- |
| Model | $df$ | $F$ | $p$ | $R^2_{Adj}$ | $df$ | $F$ | $p$ | $R^2_{Adj}$ |
| $N$ Features | (4,5911) | 1757.44 | <.0001 | 0.543 | (4,3877) | 1897.80 | <.0001 | 0.662 |
| $N$ Features + Classifier | (6,5909) | 1260.82 | <.0001 | 0.561 | (5,3876) | 1592.54 | <.0001 | 0.672 |
| $N$ Features + Classifier + Method | (12,5903) | 2550.96 | <.0001 | 0.838 | (11,3870) | 2486.92 | <.0001 | 0.876 |
| $N$ Features + Classifier + Method + Interaction Terms | (24,5891) | 1356.07 | <.0001 | 0.846 | (17,3864) | 1621.99 | <.0001 | 0.877 |

**Table 12.** Final Model Regression Results without the SVC classifier for HCP.

| Parameter Subset | UCLA |  |  |  | HCP |  |  |  |
| --- | --- | --- | --- | --- | --- | --- | --- | --- |
| Parameter | $b$ (SE) | $t$ | $p$ | $\eta^2_{part}$ | $B$ (SE) | $t$ | $p$ | $\eta^2_{part}$ |
| Feature Set Size |  |  |  |  |  |  |  |  |
| *N Features* | <0.001<br>(<0.001) | 14.87 | <0.001 | 0.422 | <0.001<br>(<0.001) | -1.24 | 0.217 | 0.316 |
| $\log_{10}(N \text{ Features})$ | 0.194<br>(0.015) | 13.13 | <0.001 | 0.717 | 0.616<br>(0.018) | 33.98 | <0.001 | 0.810 |
| $\sqrt{(N \text{ Features})}$ | -0.024<br>(0.003) | -9.45 | <0.001 | 0.209 | 0.024<br>(0.003) | 7.71 | <0.001 | 0.387 |
| $\sqrt[3]{(N \text{ Features})}$ | 0.073<br>(0.012) | 5.94 | <0.001 | 0.018 | -0.189<br>(0.015) | -12.42 | <0.001 | 0.031 |
| Classifier |  |  |  |  |  |  |  |  |
| Random Forest (RF) | -0.002<br>(0.003) | -0.91 | 0.362 | 0.072 | - | - | - | - |
| Ridge Regression (RR) | 0.034<br>(0.003) | 12.61 | <0.001 | 0.039 | 0.033<br>(0.003) | 12.09 | <0.001 | 0.080 |
| Method |  |  |  |  |  |  |  |  |
| HC | -0.002<br>(0.007) | -0.34 | 0.732 | 0.042 | 0.03<br>(0.007) | 4.07 | <0.001 | 0.004 |
| SFM | 0.096<br>(0.006) | 15.30 | <0.001 | 0.010 | 0.088<br>(0.006) | 14.14 | <0.001 | 0.009 |
| RFPI | 0.177<br>(0.006) | 28.17 | <0.001 | 0.137 | 0.116<br>(0.006) | 18.61 | <0.001 | 0.039 |
| kPCA | 0.245<br>(0.005) | 50.11 | <0.001 | 0.399 | 0.2<br>(0.005) | 41.20 | <0.001 | 0.411 |
| PCA | 0.244<br>(0.007) | 36.77 | <0.001 | 0.302 | 0.199<br>(0.007) | 30.23 | <0.001 | 0.302 |
| tSVD | 0.208<br>(0.006) | 34.26 | <0.001 | 0.332 | 0.165<br>(0.006) | 27.38 | <0.001 | 0.321 |
| Classifier*Method |  |  |  |  |  |  |  |  |
| RF*HC | 0.004<br>(0.009) | 0.38 | 0.706 | <0.001 | - | - | - | - |
| RF*SFM | -0.01<br>(0.009) | -1.16 | 0.245 | <0.001 | - | - | - | - |
| RF*RFPI | -0.023<br>(0.009) | -2.55 | 0.011 | <0.001 | - | - | - | - |
| RF*kPCA | -0.086<br>(0.007) | -12.41 | <0.001 | 0.019 | - | - | - | - |
| RF*PCA | -0.077<br>(0.009) | -8.17 | <0.001 | 0.009 | - | - | - | - |
| RF*tSVD | -0.073<br>(0.009) | -8.47 | <0.001 | 0.020 | - | - | - | - |
| RR*HC | 0.015<br>(0.009) | 1.61 | 0.107 | <0.001 | 0.021<br>(0.01) | 2.06 | 0.040 | <0.001 |
| RR*SFM | -0.004<br>(0.009) | -0.49 | 0.622 | <0.001 | -0.002<br>(0.009) | -0.25 | 0.802 | <0.001 |
| RR*RFPI | -0.012<br>(0.009) | -1.32 | 0.187 | <0.001 | -0.012<br>(0.009) | -1.40 | 0.161 | 0.001 |
| RR*kPCA | -0.018<br>(0.007) | -2.68 | 0.007 | 0.001 | 0.02<br>(0.007) | 2.90 | 0.004 | 0.001 |
| RR*PCA | -0.017<br>(0.009) | -1.81 | 0.070 | <0.001 | 0.018<br>(0.009) | 1.95 | 0.052 | <0.001 |
| RR*tSVD | 0.019<br>(0.009) | 2.26 | 0.024 | <0.001 | 0.035<br>(0.009) | 4.07 | <0.001 | 0.004 |

*Note:* HC Hierarchical Clustering, RFPI: Random Forest Permutation Importance, SFM: Select From Model, tSVD: Truncated Singular Value Decomposition, PCA: Principal Component Analysis, kPCA: Kernel Principal Component Analysis, LDA: Linear Discriminant Analysis, RR: Ridge Regression, RF: Random Forest, SVC: Support Vector Classifier

## 4. Discussion

The present study examined the effects of utilizing a suite of DR and FS methods on cognitive task decoding in two publicly available fMRI datasets. At a broad level, both DR and FS methods performed better than simply providing the classifiers the full input data (19,990 features) or by randomly selecting features. This suggests that such methods can capture meaningful information in functional connectivity features regarding cognitive task decoding. Figures 1 and 2 demonstrate the overall trends of each method and classifier with respect to the full number of possible features selected, from small to large. In several configurations of classifier and DR/FS method, an inverted parabolic function in the number of features is noted, where the optimal decoding accuracy is between roughly 100-2,000 features. These methods often perform better than randomly selecting features, suggesting that useful information is contained in the specific smaller number of features selected, or combined through a DR approach. This “sweet spot” for features has pragmatic computational implications. Considering a baseline total feature set of 19,990 features, this suggests that optimal classification performance can be achieved with many of these methods with only .005-.10% of the total feature space. Thus, a large reduction in the total computational complexity of predictive models can result in very good classification performance.

Higher accuracy was obtained in the HCP dataset relative to the UCLA dataset across methods. Some potential reasons for this discrepancy may be scanning parameters, data quality, or the nature of the tasks themselves. However, the support vector machine had poor performance in the HCP dataset overall. Task-level analysis demonstrated that the SVC in HCP in the top 10% of performing SVCs suggested that certain tasks were more difficult to decode for the SVC. This difficulty was not replicated in the other classifiers, nor was this effect demonstrated in the UCLA dataset. In the UCLA dataset, some tasks were not decoded at a rate better than baseline methods in the top 10% of performing models. However, this was not the case in HCP, where there was better task-level decoding. Additionally, in the UCLA dataset, there was more variability in the decoding of tasks, where some tasks, such as the breath-holding tasks, were classified more accurately relative to other tasks. Given the overall greater degree of decoding accuracy in HCP, these results suggest caution in the selection of classifiers in future research, such that certain datasets may have certain characteristics that will cause some classifiers to reach failure modes.

When considering all the classifiers and DR/FS methods, our hierarchical regression approach found that DR/FS approaches factored into greater classifier performance in the UCLA than HCP dataset. However, when the poorly performing SVC was removed from the model, this difference disappeared. In both datasets, DR/FS methods accounted for a large Δ*R*^2^, supporting the idea that DR/FS method matters above and beyond classifier and feature set size.

While the present study focuses on decoding accuracy as the optimal outcome, decoding accuracy may not be the sole metric to optimize. A relative advantage of the classifiers used in the present study, and by extension FS methods relative to DR approaches, is the fact that they are more interpretable. In the context of more complex deep learning approaches, which can often be “black box”, these methods can identify particular features associated with the outcome variable of interest. In the HCP dataset in particular, rates of decoding accuracy were often over 95%. The use of more complex deep learning models would not offer substantive gains in accuracy and would be at the expense of interpretability (Rudin, 2019). As an example, the decoding accuracy obtained in the present study in the HCP dataset exceeds that of recent research utilizing more complex modeling architectures (Nishimura et al., 2024). Additionally, the usage of simpler classifiers like the ones in the present study allow for large scale hyperparameter tuning and cross validation, which may not be tractable with deep learning models. In addition to interpretability, some researchers have argued that environmental impact should be considered part of what makes a classifier the “best” performing (Schwartz et al., 2019). In Figure 3, we plot the runtime of each set of features for each of the different classifiers. This visually represents the time it takes for each classifier with each DR/FS method to be run through the entire cross validated training and testing process. At around 1,000 features, the duration of the runtime dramatically increases. However, this threshold falls within the identified window of higher decoding accuracies in figures 1 and 2. This suggests that an optimal tradeoff between runtime and decoding accuracy can be achieved. This effect was noted in both HCP and UCLA datasets. It is possible that this tradeoff can inform future DR/FS implementations with other fMRI decoding datasets, especially as the energy footprint of developing, training, and testing models will grow to be a larger consideration of ML research in neuroscience.

**Figure 3.**
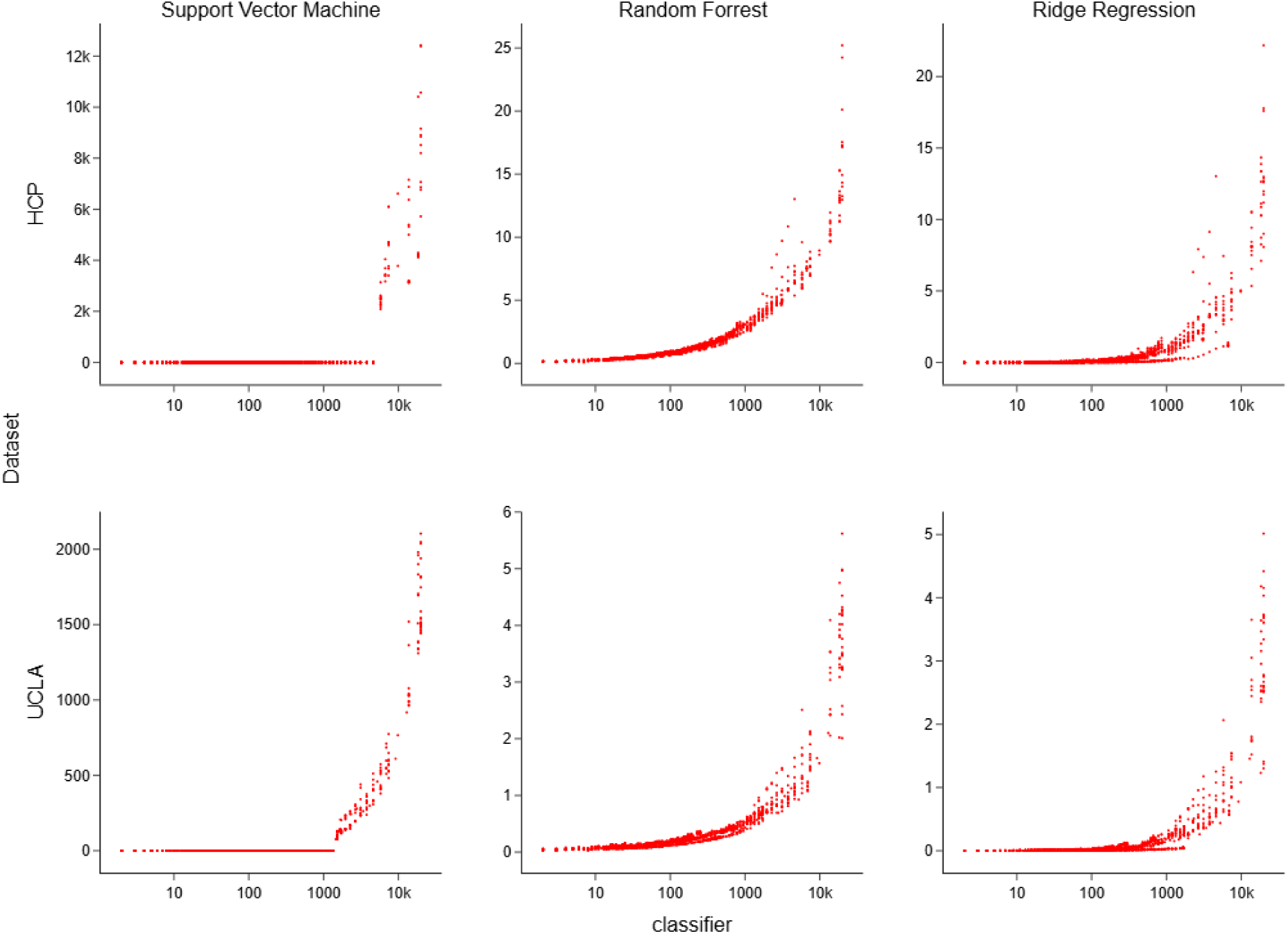
Classifier runtime as a function of feature sizes and dataset.

Our task-level analysis suggests that some DR/FS methods and classifiers were better at decoding certain tasks relative to others. However, some tasks, in particular the social processing task in HCP, had a high degree of decoding accuracy regardless of classifier or DR/FS method. However, more variance was noted in the UCLA dataset, where the breath holding task (which was the most accurately decoded task) was not as consistently classified across all DR/FS methods. This suggests that some cognitive tasks have high levels of signal-to-noise ratios relative to other tasks. These results pose several potentially exciting future directions regarding the decodable information contained in task-based fMRI. In particular, it will be useful to examine what makes some tasks robust to methods for a high level of decoding accuracy across classifiers. Potentially, task design or cognitive mechanisms might explain why some tasks are more decodable than others. Future research can perhaps replicate these task designs to better examine factors that contribute to their decodability.

### 4.1 Limitations

A key advantage of this study is the reliance upon methods that can be fit on a separate sample of data than the test set. While many DR or FS methods can be applied regardless of dataset, applying methods that can be tested on data not in the original DR or FS step prevents the possibility of any data leakage from the test set in evaluation of classification accuracy. Leakage of information from the training set into the testing set can be a source of inflated estimates of accuracy (Rosenblatt et al., 2023). The usage of these methods that can be fit on training data and applied to testing data provide unbiased estimates of their performance.

We selected methods that operated in this fashion in the present study. Many dimensionality reduction methods exist that do not have the same ability as the ones we have employed here. Thus, while we acknowledge that the methods used in the study are not comprehensive, they utilize best practices in machine learning to reduce the likelihood of overfitting, leakage, and other methodological problems that can cause inflated estimates of accuracy, which can be a problem in the machine learning and brain imaging literature (Varoquax et al., 2017). Additionally, the number of potential classifiers to choose from are quite large, especially with respect to traditional machine learning classifiers. Our choice of classifiers was motivated by frequently employed methods currently used in the fMRI literature that were not wrapped methods that conduct inherent DR or FS (i.e., LASSO or elastic net regression). Additionally, given the large number of classifiers with a large number of DR and FS methods being subject to a grid search for optimal hyperparameters and cross validation, some classifiers (e.g., XGBoost, k-nearest neighbors) can have longer run times at scale that make them prohibitively costly to test at the scope that were tested in the present study (Bhatia, 2010; Bentéjac et al., 2021).

### 4.2 Future Directions

The results of the present study suggest that in some datasets, near-perfect decoding accuracy can be achieved using traditional machine learning classifiers with DR and FS methods. The present study utilized static functional connectivity for decoding. While static connectivity is relatively lower in terms of dimensionality, method comparisons such as the ones performed in the present study should be undertaken with dynamic functional connectivity (Spencer & Goodfellow, 2022) or other imaging modalities such as electroencephalography (EEG; García-Laencina et al., 2014). Future research should also examine the potential drivers of the “peak” in performance around 100-2,000 features. This work could explore whether these results hold with other methods as well as other datasets. These findings can then inform future decoding research. It would also be of note to explore whether this effect holds of a certain peak of features results in optimal classification in fMRI research outside of cognitive task decoding schemes. Additionally, the context of decoding in the present study was the overall task level. Future research could examine whether this decoding holds at either the block or the trial level within a task, or across broad types of tasks. While each study contained a working memory task, they are difficult to directly compare due to differences in the type of contents in working memory. More finely grained cognitive task phenotyping could be a useful methodological approach to the type of comparative data analysis utilized in the present study (Poldrack and Yarkoni, 2016). Another potential avenue is to explore whether these methods may benefit the decoding of phenotypic information, such as age, sex, psychiatric disorders, or neurological diseases. Finally, while test and train splits were utilized to test the fit of the DR and FS methods utilized in the present study, future work should combine samples of data together where possible to examine if DR or FS methods may still be effective in identical tasks collected in different samples, which can elucidate the contribution of differences in scanner and other protocol steps involved in neuroimaging research.

### 4.3 Conclusion

The present study examined the effectiveness of various DR and FS methods in the context of cognitive task decoding in two open source datasets. These methods performed better than baseline methods, suggesting that DR/FS help improve the signal to noise ratio with respect to decoding. At the dataset level, neither DR or FS appeared to be superior to one another. However, specific DR and FS methods varied in effectiveness with type of task being decoded, with some methods not performing better than baseline for some tasks.

An important advantage of FS over DR methods is that features used for classification can be identified. Trends in the size of features were also evident. The number of features can be reduced by a large degree to still achieve at times near-perfect decoding accuracy using traditional machine learning models. This can be of benefit to researchers who are interested in neuroimaging research but may not have access to large scale computing resources. Furthermore, this accuracy can be achieved at low computational cost, thereby demonstrating a route forward for optimizing decoding performance while also minimizing environmental impact. The current trend in neuroimaging is for the size of datasets to increase. As the volume of neuroimaging data and tasks in decoding analysis grows, these findings will have implications for research or design of brain machine interfaces that maximize predictive accuracy while accounting for a large number of predictor variables and while minimizing computational and environmental cost.

## Acknowledgments

We would like to thank Greg Miller for his support providing computational resources to conduct the analyses in the present paper.

## Code

Scripts used to generate the results in this manuscript are available at https://github.com/coreyjr2/HCP-Analyses.

